# Exercise does not enhance short-term deprivation-induced ocular dominance plasticity: evidence from dichoptic surround suppression

**DOI:** 10.1101/2020.10.07.329896

**Authors:** Alex S Baldwin, Hayden M Green, Abigail E Finn, Nicholas Gant, Robert F Hess

## Abstract

The input from the two eyes is combined in the brain. In this combination, the relative strength of the input from each eye is determined by the ocular dominance. Recent work has shown that this dominance can be temporarily shifted. Covering one eye with an eye patch for a few hours makes its contribution stronger. It has been proposed that this shift can be enhanced by exercise. Here, we test this hypothesis using a dichoptic surround suppression task, and with exercise performed according to American College of Sport Medicine guidelines. We measured detection thresholds for patches of sinusoidal grating shown to one eye. When an annular mask grating was shown simultaneously to the other eye, thresholds were elevated. The difference in the elevation found in each eye is our measure of relative eye dominance. We made these measurements before and after 120 minutes of monocular deprivation (with an eye patch). In the control condition, subjects rested during this time. For the exercise condition, 30 minutes of exercise were performed at the beginning of the patching period. This was followed by 90 minutes of rest. We find that patching results in a shift in ocular dominance that can be measured using dichoptic surround suppression. However, we find no effect of exercise on the magnitude of this shift. We further performed a meta-analysis on the four studies that have examined the effects of exercise on the dominance shift. Looking across these studies, we find no evidence for such an effect.

## 1 Introduction

In binocular vision, the inputs from the two eyes are combined in the primary visual cortex. The relative weight given to the input from one eye over the other is called the “ocular dominance”. Physiologists have revealed maps of areas of the cortex where the two eye inputs are combined with different weights (Hubel and Wiesel, 1962, 1969; Shmuel et al., 2010). These “ocular dominance columns” divide the cortex into areas favouring either eye. At a higher level however, behavioural studies can also measure an overall ocular dominance. They do so with tasks where the relative combination or competition between the two eyes can be measured (Miles, 1930; Coren and Kaplan, 1973; Ding et al., 2018). Recent studies have shown that the ocular dominance can be altered. Covering one eye with a patch for a short period (e.g. two hours) results in a relative increase in that eye’s contribution to binocular vision (Lunghi et al., 2011). This shift is transient, with the ocular dominance returning to the baseline over a period of approximately one hour. This “ocular dominance plasticity” effect was first demonstrated using a binocular rivalry task (Lunghi et al., 2011). It was then confirmed to also affect a series of binocular combination tasks (Zhou et al., 2013; Spiegel et al., 2017). Imaging studies show effects of the deprivation in primary visual cortex (Tso et al., 2017; Chadnova et al., 2017; Zhou et al., 2015; Binda et al., 2018). There is evidence from magnetic resonance spectroscopy that the shift involves a GABA-ergic modulation (Lunghi et al., 2015).

Recently, it has been shown that short term patching of one eye also modulates dichoptic surround suppression (Serrano-Pedraza et al., 2015). In general, surround suppression occurs when the response to a stimulus is reduced due to the presence of other nearby stimuli. In some contexts the opposite effect can occur (surround facilitation). Such surround interactions are ubiquitous in vision. They occur at all stages of the visual pathway from the retina (McIlwain, 1964; Solomon et al., 2006), through the LGN (Levick et al., 1972; Marrocco et al., 1982; Bonin et al., 2005; Sceniak et al., 2006; Alitto and Usrey, 2008), to the primary visual cortex (Hubel and Wiesel, 1965; Blakemore and Tobin, 1972; Maffei and Fiorentini, 1976; Gilbert, 1977; Nelson and Frost, 1978; Sceniak et al., 2001; Cavanaugh et al., 2002; Van den Bergh et al., 2010; Angelucci and Shushruth, 2013) and extrastriate cortex (Allman et al., 1985; Desimone and Schein, 1987; Born and Bradley, 2005). Surround effects are also found behaviourally. The presence of a surround affects performance for threshold detection of localized stimuli (Falkner et al., 2010; Reynolds and Heeger, 2009; Sanayei et al., 2015; Foley, 2019). They also affect scene segmentation (Hupé et al., 1998; Park and Tadin, 2014) and the suprathreshold appearance of stimuli (Andriessen and Bouma, 1976; Cannon and Fullenkamp, 1991; Petrov et al., 2005; Snowden and Hammett, 1998; Zenger-Landolt and Heeger, 2003). Although surround interactions can be facilitative or suppressive, in most cases they are suppressive. Their mechanism is thought to involve the GABA neurotransmitter (Alitto and Dan, 2010; Angelucci and Bressloff, 2006; Gieselmann and Thiele, 2008; Nurminen and Angelucci, 2014; Smith, 2006). A special case of surround suppression occurs with dichoptic surrounds (where the target and surround are presented to different eyes). Depending on their methods, studies on dichoptic surround suppression have found smaller (Chubb et al., 1989) or larger effects (Meese and Hess, 2004; Petrov and McKee, 2006). These can be reconciled if dichoptic surround suppression involves separate processes occurring at different stages of the visual system (Webb et al., 2005; Cai et al., 2008, 2012; Schallmo and Murray, 2016; Schallmo et al., 2019). The recent results from Serrano-Pedraza et al. (2015) indicate that short-term patching of one eye affects the later, cortical stage of dichoptic surround suppression. Following two and a half hours of patching, the suppressive effect of a dichoptic surround presented to the non-deprived eye was reduced.

Exercise has been shown to be a powerful modulator of visual cortical plasticity in rodents (Sale et al., 2014). A recent study suggested that this is also the case in human adults (Lunghi and Sale, 2015). Lunghi and Sale (2015) performed an ocular dominance plasticity study using a binocular rivalry to measure eye dominance. They found that exercise during the deprivation period enhanced the plastic shift in eye dominance. If true, this would be the first indication that human visual plasticity can be enhanced by exercise. This would agree with previous evidence from mouse studies (Baroncelli et al., 2012). Such effects may be mediated via a common change in GABA-ergic inhibition. Since the Lunghi and Sale (2015) publication, two further studies have been published that do not confirm their conclusion. The first used a binocular combination paradigm with a comparable exercise regime (Zhou et al., 2017). The second study was an unsuccessful attempt to replicate the original result of Lunghi and Sale (2015) using a binocular rivalry task similar to theirs (Finn et al., 2019). In this study, we set out to further investigate the possible role of exercise in cortical plasticity. We tested whether exercise could enhance the effect of short-term patching on dichoptic surround suppression. This method allowed us to separately measure the effective suppression from each eye. We found no significant effect of exercise on the ocular dominance shift from short-term monocular patching. Furthermore, the patching effect we do find is restricted to a reduction in suppression of the deprived eye. This agrees with the results from the previous work by Serrano-Pedraza et al. (2015). As in their study, we do not find a reciprocal *increase* in suppression of the non-deprived eye. To date, studies examining this phenomenon have used intermittent exercise protocols of variable intensity. The relative exercise load (intensity and duration) is a key determinate of physiological responses and adaptations to exercise. In the present study we aimed to increase the ecological validity of existing findings by using an exercise dose that is recommended by health professionals and that was standardized according to participants’ individual levels of cardiorespiratory fitness.

## 2 Experimental Procedures

### 2.1 Participants

Twenty healthy adult participants volunteered to take part in this study. There were eleven female and nine male participants, with an average age of 23 years old (range 20 - 28). The average stature was 1.7 ± 0.1 metres, and average body mass was 73 ± 15 kilograms. Volunteers were eligible to participate if they were free of cardiovascular disease and musculoskeletal injury. Subjects had normal or corrected-to-normal vision. Each subject took part in three sessions after giving written informed consent. The study protocol was approved by the University of Auckland Human Participants Ethics Committee.

### 2.2 Physiological Measurements

The study protocol began with a familiarization session. A cardiopulmonary fitness test was conducted on an electromagnetically-braked cycle ergometer (Velotron Dynafit Pro, Seattle, WA, USA). We simultaneously measured ventilation and analysed expired gas composition (pneumotachometer, MLT1000L, gas analyzer, ML206, AD Instruments) to determine peak oxygen uptake (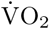 peak). Peak and submaximal oxygen uptake values were used to prescribe a workload equivalent to 60% of 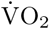 max for the Exercise condition. Heart rate was used as a secondary assessment of workload prescription. It was measured during the 2 hours of monocular deprivation using a chest strap heart rate monitor (Polar FT1, Polar Electro, Finland). Data were collected upon eye patching. Further measurements were made every five minutes in the first forty minutes of monocular deprivation. After this point measurements were made every ten minutes.

### 2.3 Psychophysical Methods

A measure of interocular suppression was obtained with a dichoptic surround suppression task. Thresholds for detecting a target grating were determined with a two-interval forced-choice design. Subjects fixated on a central marker, and a target would appear in only one eye (monocularly) either to the left or right of that marker for 300 ms. The task set to the subject was to respond whether the target appeared to the left or right of fixation. The target was a circular patch of 4 c/deg grating presented 1.5 degrees of visual angle from the fixation marker (Figure 1A). The edges of the grating were softened with a raised-cosine envelope that declined from the plateau to zero contrast over 0.25 degrees. The central plateau was 0.25 degrees wide. Therefore, the full-width at half-magnitude of the grating was 0.5 degrees. This gave approximately two visible cycles of the carrier grating within the target. In the detection threshold condition (without a dichoptic mask) the two potential target locations were surrounded by a pair of black circles during stimulus presentation (25% contrast, diameter of 1.7 deg). In the non-target eye, only the two black circles were shown (Figure 1B).

**Figure 1:**
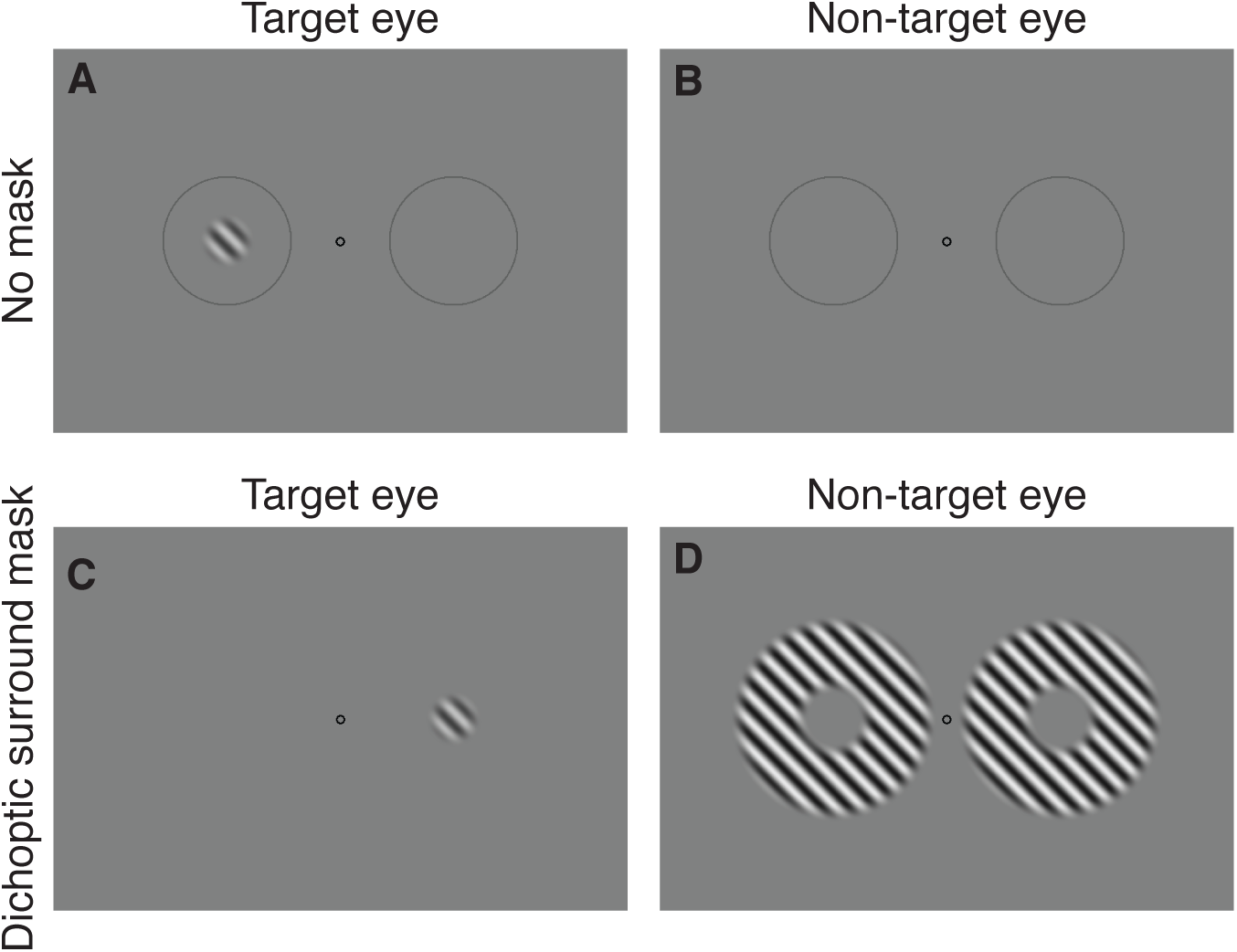
Examples of the two stimulus conditions used in this study. The top row (A-B) shows the simple detection condition where the subject detects a small patch of grating seen only by the target eye. The subject fixates the small central circle. There are two larger circles that indicate the two potential stimulus locations in this two-alternative forced-choice task. The bottom row (C-D) shows the dichoptic surround suppression condition. The stimulus presented to the target eye is the same, except the circles indicating the location of the target are removed. In the non-target eye two annular grating masks are presented, enclosing the two potential target locations.

The condition with the dichoptic surround mask is shown in Figure 1C-D. In this condition, the stimulus presented to the target eye was almost identical to that in the threshold condition. The only difference was that the two black circles surrounding the target location were not shown (Figure 1C). Instead, an annular grating was presented at both locations in the non-target eye (Figure 1D). This grating had the same spatial frequency as the target (4 c/deg), but was presented in the opposite spatial phase (a white bar in the target grating was aligned with a black bar in the annular surround grating). The same raised-cosine smoothing was applied to the surround as was applied to the target. There was also small gap between the outer edge of the target and the inner edge of the annular surround grating. At half-magnitude, the radius of the inner edge of the annular grating was 0.23 deg, and the outer edge was at 0.63 deg. The dichoptic surround mask grating was presented at 45% contrast. The target contrast was controlled by a pair of interleaved staircases (Baldwin, 2019) for each eye. Each eye had one 3-down-1-up staircase starting at 21 dB contrast and one 2-down-1-up staircase starting at 27 dB contrast. The staircase step size was 6 dB before the first reversal and 3 dB thereafter.

### 2.4 Experiment Design

Subjects first underwent a familiarisation session. This involved practicing the dichoptic surround suppression task and measurement of cardiorespiratory fitness. Fitness was measured using the maximal aerobic capacity (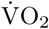 max) test. This familiarisation session was followed by two experimental sessions (see Figure 2). These began with baseline dichoptic suppression threshold measurements. Subjects then underwent two hours of monocular deprivation (MD) with an orthoptic eye patch (Nexcare, Opticlude, 3M, Canada). Following patch removal, further post-patching measures were made at 15 minute intervals (0, 15, 30, 45 minutes). In one of the experimental sessions, participants performed continuous aerobic exercise for 30 minutes. They exercised on a cycle ergometer at a moderate intensity (60% of their 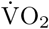 max) to achieve, at least, the minimum duration and intensity of exercise recommended by the American College of Sports Medicine for health and fitness (American College of Sports Medicine, 2017). This was followed by seated rest for the remaining 90 minutes of monocular deprivation. In both sessions, participants watched movies on a television screen throughout the 120 minutes of monocular deprivation. The order of the two experimental treatments, and the eye to be patched, were allocated randomly in a counterbalanced fashion.

**Figure 2:**
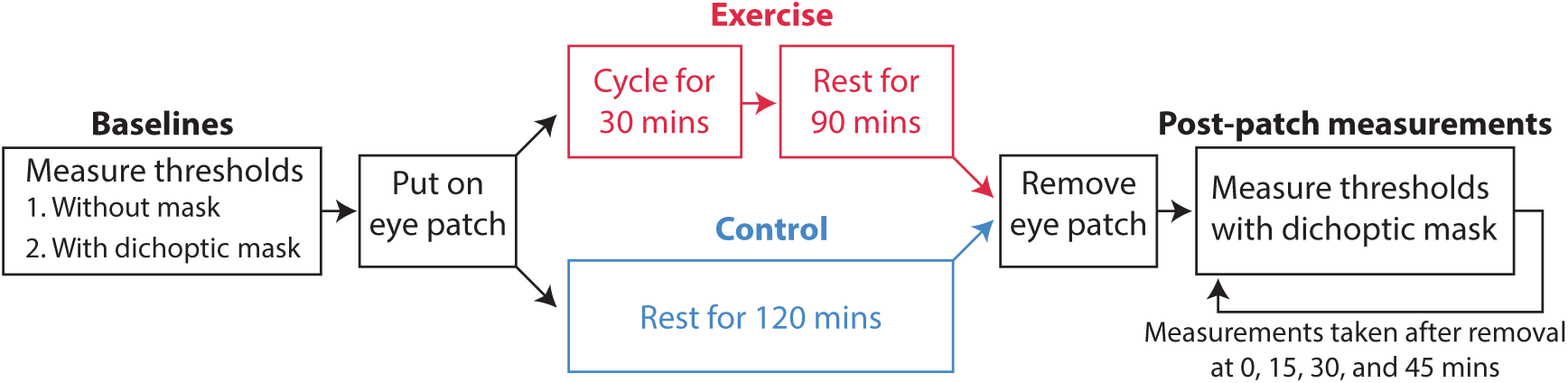
Procedure for testing. On each of the two testing days the subject performed either the exercise or the control condition. During the cycle period, subjects were watching movies on a television screen, whilst in the rest period, for both conditions, subjects sat in a chair and watched movies on a television screen.

### 2.5 Mood Ratings

Visual analogue scales (VAS) were used to subjectively assess day-to-day lifestyle changes that could influence experimental outcomes. Upon arrival to the laboratory, participants were asked to rate their motivation for the testing session, general stress levels, and diet quality over the preceding 24 hours on a VAS rated from “very bad” to “very good” for motivation and diet, and “very low” to “very high” for stress levels. During each experimental session VAS were used at four timepoints to assess valence, arousal and fatigue. These occurred before and after the baseline visual tests, after the 2 hours patching and following the last visual test. Subjects’ responses were recorded as a fraction of the scale length. Written cues for valence were “neutral” at 50% of the scale, and “very bad” and “very good” at opposite ends of the scale. Cues for arousal and fatigue were “not at all” (fatigued or aroused) and “highly fatigued/aroused” on either end of the scale.

### 2.6 Analysis

Repeated measures analyses of variance (ANOVA) were used to test the differences between experimental conditions for heart rate and subjective measures. For heart rate, valence, motivation and arousal the ANOVA factors were condition (Rest or Exercise) and time. For motivation, diet and stress levels a paired students t-test was used to compare the two conditions. In cases where assumptions of sphericity were violated Greenhouse-Geisser corrections were applied. If a main effect was deemed to be statistically significant (p < 0.05) post-hoc comparisons were adjusted using the Bonferroni correction to control for multiple comparisons. These statistical analyses were conducted in SPSS version 24 (IBM, USA).

For the psychophysical measurements, contrast detection thresholds were obtained through psychometric function fitting. The Palamedes toolbox (Prins and Kingdom, 2009) was used to perform this analysis. Data were fit with a Quick psychometric function (Quick, 1974) by a maximum-likelihood procedure. We calculate differences in threshold using dB logarithmic units. The formula is

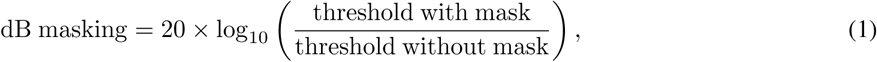

therefore, a difference of 6 dB indicates that the mask approximately doubles the contrast required for detection.

## 3 Results

### 3.1 Physiological Measurements

The heart rate measurements are presented in Figure 3. The mean 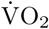 peak was 43 ± 9 ml.kg^−1^.min^−1^. The average workload performed for 30 minutes during the Exercise condition was 123 ± 34 W (equivalent to 60% of VO2 max). A technical error occurred during sub-maximal stages for one subject. Their workload was estimated using the peak workload recorded during the test. An ANOVA was performed on the heart rate measurements. There were significant main effects of condition and time as well as a condition x time interaction effect observed with the ANOVA for both main and interaction effects. As expected, post-hoc comparisons showed a higher heart rate at all time levels during physical exercise and the initial phase of the recovery period (p < 0.05 at 5-50 minutes and 80 minutes patched). Three subjects’ heart rate data were excluded from analysis due to equipment-related data loss.

**Figure 3:**
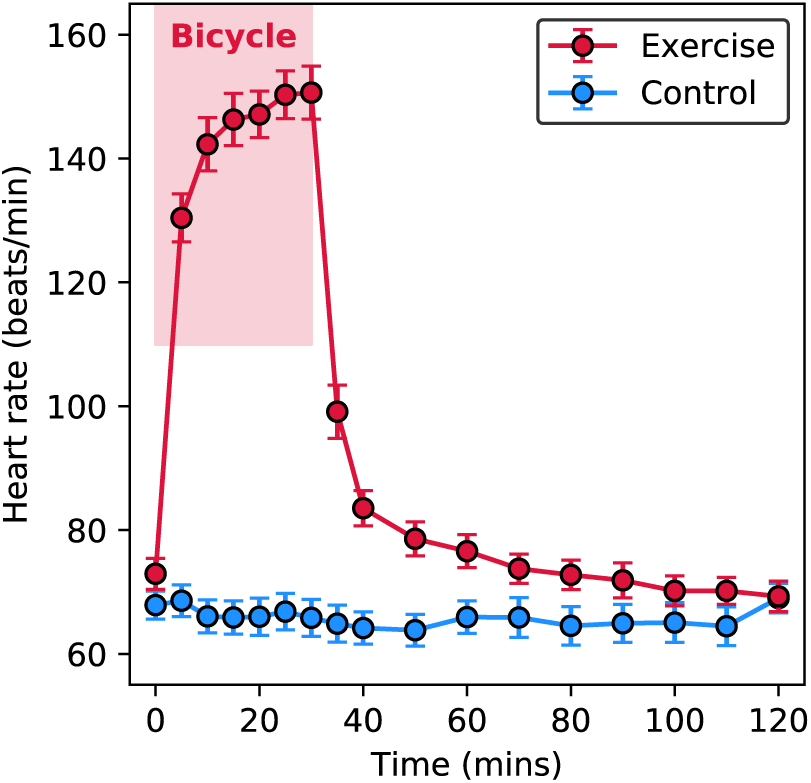
Heart Rate during the two hours of monocular deprivation for the Rest and Exercise conditions. Rest vs. Exercise p < 0.05 at time levels 5, 10, 15, 20, 25, 30, 35, 40, 50, 70 minutes.

### 3.2 Psychophysical Results

The results from the psychophysical task are presented in Figure 4. The graphs in the top row plot the threshold elevation caused by the dichoptic surround mask for each timepoint. This is given in logarithmic dB units, so that a value of 0 dB would mean “no effect”. Each 6 dB increase on this axis indicates a doubling of the unmasked threshold. The leftmost points in each plot give the baseline measurements made before patching. For the condition where the target was shown to the patched eye (Figure 4A) the average baseline measurements were between 9 and 12 dB. When the target was shown to the non-patched eye, the baselines were also around 12 dB. These masking measurements mean that the threshold contrast for detecting the target increased by around a factor of four when the surround mask (at 45% contrast) was presented in the other eye.

**Figure 4:**
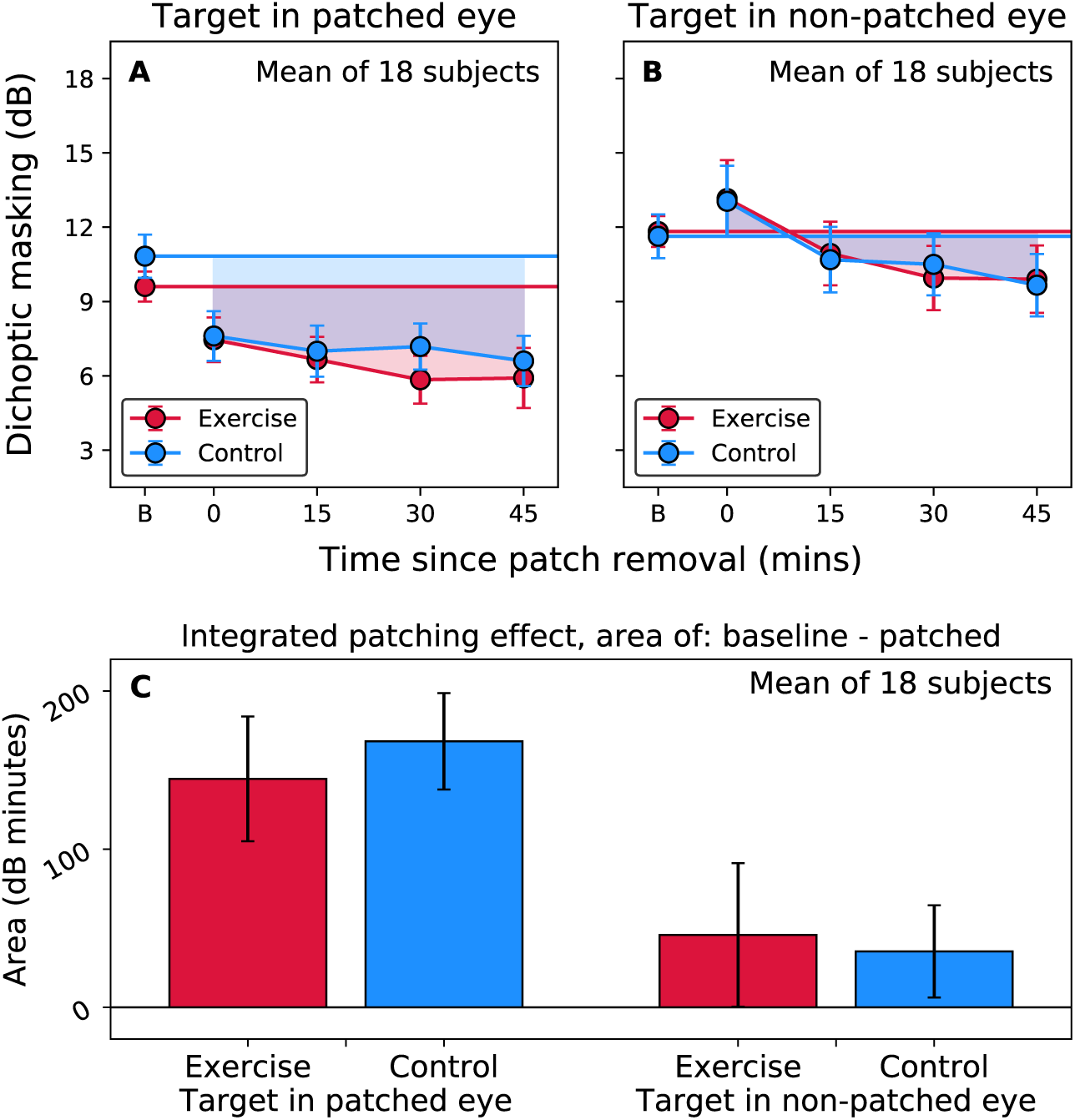
Masking from the dichoptic surround grating. Error bars give the standard error of the mean calculated over 18 subjects. Masking is presented for the condition where the target was presented to the patched eye and the mask to the non-patched eye (panel A), and vice-versa (panel B). The horizontal lines give the baseline masking, with the points at 0, 15, 30, and 45 minutes giving the masking measured at that time after patch removal. The area between the patched data and the baseline give an index of the overall patching effect (note that in panel B the 0 minute datapoint is above the baseline, giving that part an area which counts as negative in our analysis). The average areas calculated across subjects are presented in panel C. For both pairs of bars, the differences between the Exercise and Control conditions were not significant when tested with the Wilcoxon signed-rank test in SciPy.

The average threshold elevations at baseline measured on each testing day (“Exercise” and “Control”) are indicated by horizontal lines in Figure 4A-B. Any changes as a result of short-term patching are judged relative to the baseline from the appropriate testing day. That allows us to investigate how the patching affects the strength of the dichoptic interactions. Immediately following patch removal (“0” on the x-axis) the amount of masking for targets in the patched eye decreased (Figure 4A), whereas in the non-patched eye it increased slightly (Figure 4B). The amount of masking measured at this timepoint in the patched eye was almost identical for the exercise and control conditions. There was however a difference in the baseline values measured on those two testing days. This means that the shift was actually larger for the control condition (where subjects rested) than it was in the exercise condition.

Surprisingly, the measurements taken at subsequent timepoints do not show a decrease in this shift. If anything, there was a trend for the change in masking to increase over the next 45 minutes. In the non-patched eye, we also see a downward trend (less masking) over that time period. This means that by the 15 minute timepoint the effect of patching appears to have reversed. The results from both eyes therefore show a trend for decreased masking over time. We hypothesise that this trend may be a separate effect from the asymmetric shift in ocular dominance that patching causes.

We performed a three-way within-subjects ANOVA in R (RStudio Team, 2016), with factors of condition (exercise or control), target eye (patched or non-patched), and timepoint post-patching. The dependent variable was the shift in masking from the baseline measured for each subject before patching. The effect of condition alone was not significant (*F*_1,17_ = 0.05, *P* = 0.828), nor were there any significant interactions involving condition. There were significant effects of target eye (*F*_1,17_ = 11.08, *P* = 0.004) and timepoint (*F*_3,51_ = 11.78, *P* < 0.001), and a significant interaction between the two (*F*_3,51_ = 5.14, *P* = 0.004). Therefore, this analysis shows our expected patching effect, but does not support any strengthening of that effect by exercise.

Figure 4C shows the overall effect of patching. This is calculated as the area of the polygon defined by the baseline (flat lines in Figure 4A-B) and the data between the 0 and 45 minute timepoints. These are the shaded areas in Figure 4A-B. Areas above the baseline are counted as negative. These will be subtracted from the total area. We calculated these areas individually for each subject. In Figure 4C we show the mean and standard error of the 18 individual subject values. Numerically, this analysis indicates a slightly stronger patching effect in the control condition. We compared the two conditions using a a Wilcoxon signed-rank test performed in SciPy (Jones et al., 2001). The difference was not significant (*w* = 60, *P* = 0.267). The difference between the two conditions in the non-patched eye was also non-significant (*w* = 83, *P* = 0.913).

With the data from Figure 4, we performed a further analysis looking at the differences between the effects measured in the patched and non-patched eyes. In Figure 5A, each data point is essentially the result of subtracting the equivalent data point in Figure 4B from that in Figure 4A. The baseline value is now the difference between the baseline masking values measured from the two eyes. The hypothesised general downward trend in masking over time is now factored-out of the analysis. We see post-patching the typical initial peak in ocular dominance shift. This is followed by a gradual return toward the baseline value. We performed a two-way within-subjects ANOVA in R, with factors of condition (exercise or control) and timepoint post-patching. The dependent variable was the difference between the masked thresholds measured in the two eyes. This was normalised to be relative to the baseline difference calculated for each subject. In agreement with the ANOVA in the previous section, the effect of condition alone was not significant *F*_1,17_ = 2.21, *P* = 0.156. There was also no significant interaction between condition and timepoint *F*_3,51_ = 0.15, *P* = 0.927. There was however a significant effect of timepoint *F*_3,51_ = 5.14, *P* < 0.004.

**Figure 5:**
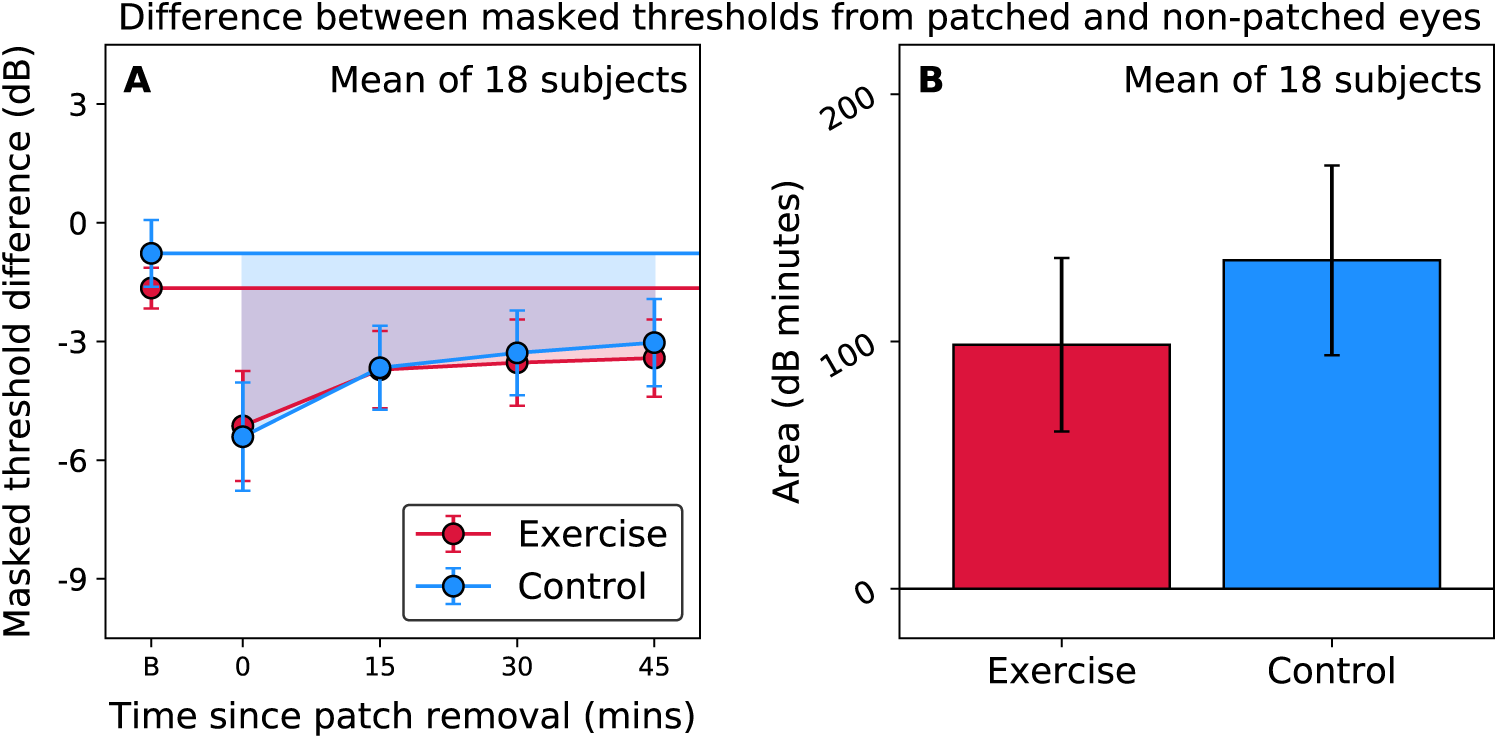
Ocular dominance analysis, based on the differences between masked thresholds (in dB units) measured from the two eyes (A). A masked threshold difference of zero dB would mean that the two eyes were balanced. The horizontal lines give the baseline imbalance, with the points at 0, 15, 30, and 45 minutes giving the measured value at each time point after patch removal. The area between the patched data and the baseline give an index of the overall patching effect. The average areas calculated across subjects are presented in panel B. The difference between the Exercise and Control conditions was not significant when tested with the Wilcoxon signed-rank test in SciPy.

We also calculated the area between the baseline and the patched data (shaded regions in Figure 5A). The mean areas calculated across 18 subjects (with standard errors) are presented in Figure 5B. As in Figure 4C, the patching effect appears to be slightly stronger under the rest condition than under the exercise condition. The direction of this trend is the opposite of the effect reported by Lunghi et al. (2015). The difference we find was not significant however by a Wilcoxon signed-rank test (w = 60, P = 0.267).

**Table 1:** Mood scores for the exercise and control conditions. Valence is rated “neutral” at 50% whilst fatigue and arousal are rated “not at all” at 0%. Values are reported as mean ± standard deviation.

|  |  | Baseline | Post-baseline | Post-MD | Post-follow-up |
| --- | --- | --- | --- | --- | --- |
| Valence (%) | Control | 80 ± 10 | 74 ± 13 | 72 ± 11 | 71 ± 14 |
| | Exercise | 79 ± 10 | 75 ± 12 | 72 ± 13 | 69 ± 14 |
| Arousal (%) | Control | 57 ± 16 | 48 ± 16 | 49 ± 15 | 42 ± 16 |
| | Exercise | 50 ± 12 | 49 ± 14 | 56 ± 11 | 43 ± 16 |
| Fatigue (%) | Control | 36 ± 19 | 37 ± 17 | 40 ± 21 | 44 ± 18 |
| | Exercise | 39 ± 21 | 41 ± 20 | 37 ± 19 | 44 ± 18 |

### 3.3 Mood and Effort Sense

There were no differences between the two conditions for motivation (*t*_18_ = 0.06, *P* = 0.956), diet (*t*_18_ = 1.38, *P* = 0.184) and stress levels (*t*_18_ = −0.56, *P* = 0.586) prior to starting the experiment. There was also no effect of condition for valence, arousal or fatigue. There was however a significant effect of time for valence (*F*_1.57,28.23_ = 12.01, *P* < 0.001), arousal (*F*_3,54_ = 10.77, *P* < 0.001), and fatigue (*F*_3,54_ = 2.99, *P* < 0.001). Valence during the experiments decreased significantly compared to baseline (*P* < 0.05). Arousal decreased following surround suppression threshold measurements that occurred before and after monocular deprivation (*P* < 0.05 baseline vs. post-baseline, post-MD vs. post-follow up and baseline vs. post follow-up) but did not decrease over the monocular deprivation period (*P* > 0.05 post-baseline vs. post-MD). One participant was excluded from the analysis due to incorrectly answered questionnaires.

## 4 Discussion

In this study we replicate the finding first reported by Serrano-Pedraza et al. (2015) that short-term monocular deprivation results in a modulation of dichoptic surround suppression. This is consistent with deprivation strengthening the binocular contribution of the patched eye (Lunghi et al., 2011; Zhou et al., 2013). When the surround mask is presented to the eye that had been patched, its suppressive effect on a target in the non-patched eye is (mildly) enhanced. When the target is presented to the patched eye and the mask to the non-patched eye, the suppression is diminished. These near surround effects are thought to involve horizontal connections in V1 (Angelucci et al., 2017). We assume that their dichoptic nature reflects interactions across eye columns in layer 4 of V1 (Yoshioka et al., 1996) or between binocular neurones in more superficial layers of V1 (Webb et al., 2005). The site of the ocular dominance changes induced by short-term deprivation is also throught to be in V1. This is based on evidence from human psychophysics (Zhou et al., 2014), and primate brain imaging (Tso et al., 2017; Chadnova et al., 2017; Lunghi et al., 2015; Binda et al., 2018).

Standardised exercise on a stationary bicycle was not shown to enhance the effect of monocular deprivation in this study. The current study is the fourth investigation of whether exercise enhances the shift in ocular dominance induced by short-term monocular deprivation. The first of these (Lunghi et al., 2015) found such an effect. This was followed-up by Zhou et al. (2017), who found no effect of exercise. A possible distinction between those two studies was that Lunghi et al. (2015) used a binocular rivalry task, whereas Zhou et al. (2017) used a phase combination task. It has been shown that the effects of monocular deprivation can vary depending on the task used to measure them (Bai et al., 2017; Baldwin and Hess, 2018). For that reason, Finn et al. (2019) attempted to replicate Lunghi et al. (2015) with a binocular rivalry task. They found no significant effect of exercise, and so failed to replicate the original finding. Previous studies have all used intermittent exercise of fixed intensity, but in the present study we increased ecological validity by prescribing exercise according to recognised guidelines and individualised capacities (determined by cardiorespiratory fitness test). We found that exercise performed according to the American College of Sport Medicine’s guidelines (American College of Sports Medicine, 2017) for maintaining physical health and well-being did not increase ocular-dominance shift. It may be inferred that an ocular-dominance shift is unlikely to be elicited by doses of physical activity undertaken by members of the exercising public following recommended guidelines.

We performed a meta-analysis by combining the effects calculated from data from four studies looking at the effects of exercise on ocular dominance plasticity (including this study). These data are summarised in Figure 6. For Lunghi et al. (2015), the ocular dominance index measures were calculated. The use of this measure (rather than the mean phase duration used in the original study) is justified in Finn et al. (2019). A modified Cohen’s d score was calculated

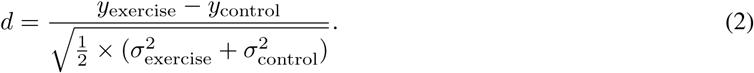

**Figure 6:**
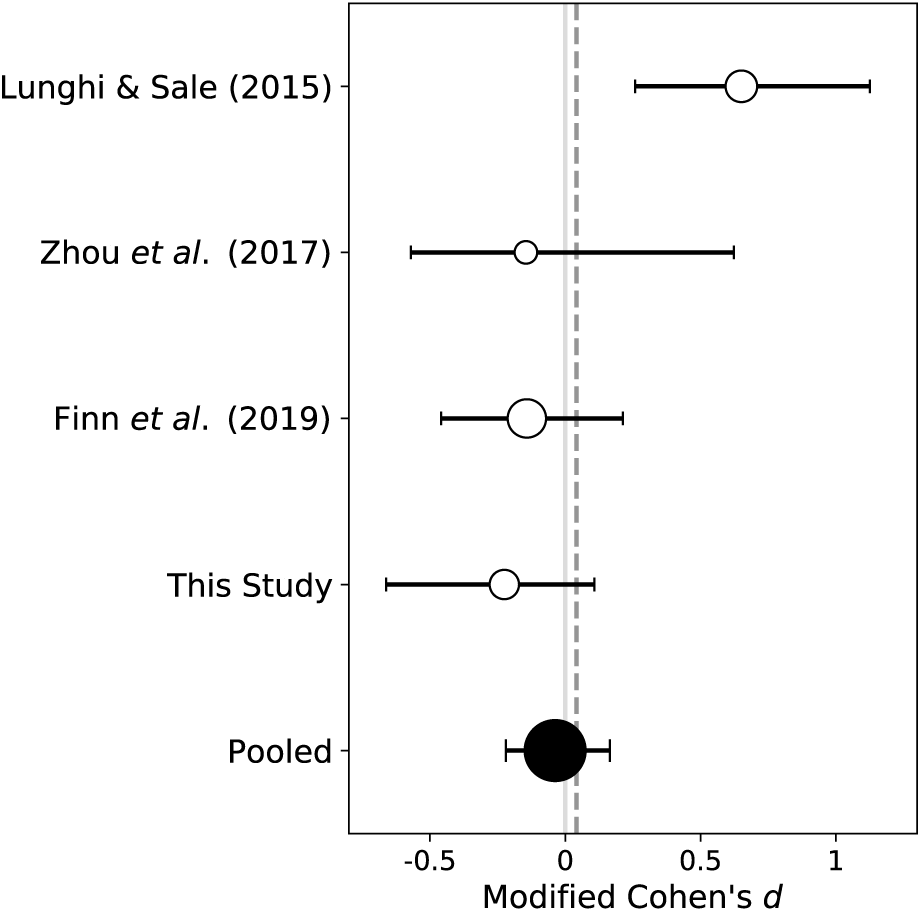
Data were taken from the four studies looking for an effect of exercise on the ocular-dominance shift. A meta-analysis was performed looking at the main result (effect of exercise) over those four studies. The areas of the circle marker symbols are proportional to the sample size in each study. The bottom symbol gives the result from pooling data across all four studies. The vertical grey line at zero indicates no effect. The dashed black line is the average of the four points, weighted by the number of subjects in each study..

The markers in Figure 6 give the values calculated from each study’s data. The bottom data point is the data pooled across all four studies. Data were normalised by the mean patching effect obtained (averaged across exercise and control) in their study before pooling. A score of zero indicates no effect of exercise. Positive numbers mean that exercise enhances the shift in ocular dominance. Negative numbers mean that exercise suppresses the shift. The error bars on the points in Figure 5 give the 95% confidence interval calculated from non-parametric bootstrapping (2000 samples). We only see evidence for an effect in the first study Lunghi et al. (2015). In fact, the three subsequent studies all show small shifts (non-significant) in the opposite direction. In a similar vein, it has recently been shown that exercise does not enhance visual perceptual learning either (Connell et al., 2018; Campana et al., 2020).

## 5 Additional information

## 5.1 Acknowledgements

This work was funded both by an ERA-NET Neuron grant (JTC 2015) awarded to Robert F. Hess, and by University of Auckland funding to Nicholas Gant. This pre-print manuscript was written in LATEX using TeXShop. It is formatted with a custom style available at: github.com/alexsbaldwin/biorxiv-inspired-latex-style.

## 5.2 Author contributions in CREDIT format

**Alex S Baldwin**: Conceptualization, Methodology, Software, Validation, Formal Analysis, Investigation, Data Curation, Writing - Original Draft, Writing - Review & Editing, Visualization, Supervision, Funding Acquisition; **Abigail E Finn**: Conceptualization, Methodology, Validation, Investigation, Writing - Review & Editing; **Hayden M Green**: Conceptualization, Methodology, Validation, Investigation, Formal Analysis, Investigation, Data Curation, Writing - Original Draft, Writing - Review & Editing, Visualization, Supervision; **Nicholas Gant**: Conceptualization, Methodology, Resources, Writing - Review & Editing, Supervision, Project Administration, Funding Acquisition; **Robert F Hess**: Conceptualization, Methodology, Resources, Writing - Review & Editing, Supervision, Project Administration, Funding Acquisition.

## 6 Appendix: outlier removal

### 6.1 Experiment Design

**Figure 7:**
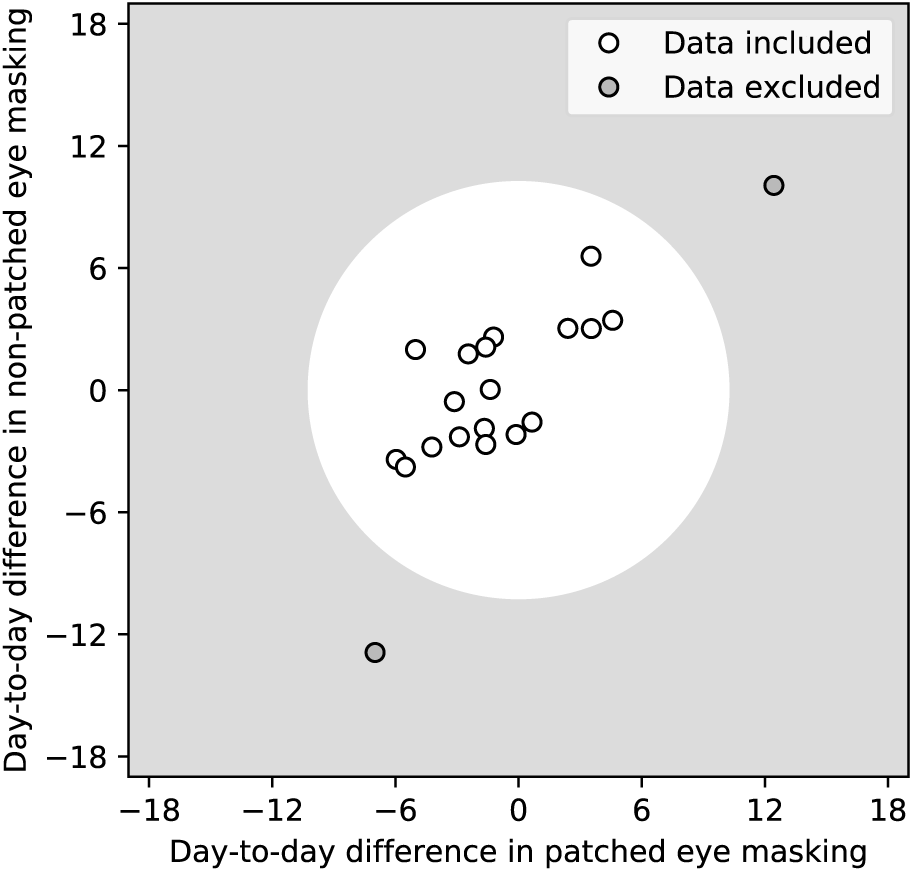
The method by which outlier subjects were removed. The day-to-day differences in the baseline masking effect measured in the patched (*x*-axis) and non-patched (*y*-axis) eyes are plotted. Data outside the white circle are outliers. The criteria for inclusion were determined using a method based on that proposed by (Tukey, 1970). The mean-normalised data were combined for the two eyes. The first (Q1) and third (Q3) quartiles were found, and the interquartile range (IQR). The maximum allowable day-to-day difference was the average of |Q1 - 1.5 × IQR| and |Q3 + 1.5 × IQR|, this came out as a 10 dB maximum day-to-day difference.

Of the twenty subjects we tested, two were removed from further analysis. The removal was based on the day-to-day differences in the dichoptic surround masking measured at baseline. The method by which this was performed is illustrated in Figure S1.

